# Structurally informed evolutionary models improve phylogenetic reconstruction for emerging, seasonal, and pandemic influenza viruses

**DOI:** 10.1101/228692

**Authors:** Xueting Qiu, Justin Bahl

## Abstract

Precise estimation of genetic substitution patterns is critical for accurate reconstruction of pathogen phylogenies. Few studies of viral evolution account for variations of mutation rate across a single gene. This is especially true when considering evolution of segmented viruses where individual segments are short, encoding for few proteins. However, the structural and functional partitions of these proteins could provide valuable information for more accurate inference of viral evolution, due to the disparate immune selection pressure on different functional domains. Accurately reconstructed evolutionary features on specific functional domains can in turn provide biological information on viral protein and immune targets for vaccine design. In this study we developed and evaluated a structurally informed partitioning scheme that accounts for rate variation among immunogenic head and stalk domains of the surface protein hemagglutinin (HA) of influenza viruses. We evaluated the model fit and performance of four different models - HKY, SRD06 codon, HKY with a structurally informed partitioning scheme, SRD06 with a structurally informed partitioning scheme on pandemic A/H1N1pdm09, seasonal A/H1N1postpdm, A/H3N2, B-Yamagata-like and Victoria-like lineages, and two highly pathogenic avian influenza A viruses H5Nx and H7N9. Results showed that structurally informed partitioning with SRD06 performed better for all datasets with decisively statistical support. Significantly faster nucleotide substitution rates for head domain, compared to stalk domain was observed and may provide insight for stalk derived broadly-reactive vaccine design. Taken together, integrating a functionally informed partitioning scheme based on protein structures of immune targets allows for significant improvement of phylogenetic analysis and providing important biological insights.

## Introduction

The importance of statistical phylogenetic methods to study the epidemiology, evolution and ecology of rapidly evolving viral pathogens has been driven by the growing availability of whole genome sequences (1). Precisely estimating the pattern of genetic variations is critical to reconstruct the accurate pathogen phylogenies (2, 3). Nucleotide substitution models have been developed to describe the process of change from one nucleic state to another among viral isolates, where they often allow for rate variations between transitions and transversions (4), or incorporate nucleotide base frequencies with substitution rate parameters (5, 6). Segmented viruses, such as influenza A virus (IAV), contain short gene segments that encode for few proteins (7). Consequently, the literature regarding model development of phylogeny reconstruction is dominated by the analysis on complete protein coding regions. For example, the complete viral surface glycoprotein hemagglutinin (HA) is used the most when studying influenza viral evolution and diffusion (8). However, assuming natural selection acts uniformly across the protein domains may not be justified (9). Some partitioning strategies have been developed to account for variations inside one gene segment (3, 10). For example, approximate codon-models, such as the SRD06, incorporate information about the genetic code by allowing for a rate variation by defining substitution models for codon positions 1 and 2 and an independent model for codon position 3. This model accounts for the rapid accumulation of synonymous mutations in the third codon position (10).

HA glycoprotein with a spike-shaped structure binds to the receptors on the targeted host cell membrane when the virus begins the infection (11). This protein has two main domains – the globular head domain and stalk domain. Functionally different, the globular head domain contains the receptor binding sites, while the stalk domain is a main structure responsible for membrane fusion machinery (11, 12). Even though immune selection strongly drives antigenic drift, allowing for accumulation of mutations in the head domain, the stalk domain is functionally conserved across viral subtypes. Current seasonal influenza virus vaccines induce humoral immune responses primarily targeting the immunodominant globular head domain (13, 14), which can provide strong protection from an infecting virus that is similar with vaccine candidates, but have weak cross-reactivity to antigenic drift variants. Conversely, studies in the ferret experimental model demonstrate that the stalk domain is highly conserved across IAV subtypes and stalk specific antibodies could provide cross-reactive protection to a high diversity of IAV subtypes (13, 15). Despite the importance of understanding how substitution rates and patterns vary across functionally conserved domains in universal vaccine design (16, 17), few studies have incorporated protein structure in phylogenetic models.

In this study we proposed a novel phylogenetic model that incorporates a protein structure informed partitioning scheme to account for variable evolutionary rates resulting from immune selection. We aimed to evaluate the appropriateness of the novel model to reliably to estimate biologically informative parameters from available genetic data. We evaluated and compared four different models: HKY, SRD06 codon, HKY with a structurally informed partitioning scheme, SRD06 with a structurally informed partitioning scheme. We also evaluated the model performance across multiple viral subtypes, including pandemic H1N1pdm09, seasonal A/H1N1postpdm, A/H3N2, the two seasonal Influenza B virus lineages – Yamagata-like and Victoria-like, and two highly pathogenic avian Influenza (HPAI) H5Nx and H7N9 viruses. We further conducted sensitivity analysis to determine whether the new model was sensitive to sample size, data distribution and epidemic stage resulting in biased estimation. Separate estimation of viral evolutionary rates for the head and stalk domain may be informative for vaccine design. We therefore formally tested the hypothesis that evolutionary rates in each domain were significantly different and further estimated the relative selection pressure for each functional domain of HA protein.

### New Approaches

This study proposed a new model to incorporate a structurally informed partitioning scheme on a single protein into phylogenetic reconstruction. Decisive statistical support from the model selection procedure and stably supported across multiple influenza subtypes validated the superiority of the new model. The model can provide biological insights for viral evolution, with reporting domain-specific evolution rates and approximate selection pressure. The tree branches with domain-specific rate ratio *d*C_1+2_/*d*C_3_ can inform the approximate selection pressure on each strain, suggesting some biological explanations related to antigenic drift and emerging strains. The structurally informed phylogenetic model may reveal novel biological insights of viral evolution and have the potential to reveal more biological realism without over-parameterization.

## Results

### Model selection

To compare the four models, the log scale Bayes Factor (BF) was used, which was the difference of log marginal likelihood estimations from two models. Marginal likelihood estimation was conducted with two approaches – path-sampling and stepping-stone sampling, where both generated consistent results. Based on the model selection procedure for all subtypes (Fig. 1), the model with mixed partitioning on both codon positions and HA protein structure (CP model in Fig. 1c) had decisive BF support for better model fitting in all datasets compared to HKY and HKY with protein structure partitioning model (HKY model and P model in Fig. 1c). Compared to SRD06 codon model (C model in Fig. 1c), CP model performed significantly better for all datasets, with different levels of BF supports. HKY with protein structure model (P model) performed significantly better than HKY model for human influenza A and B viruses, but it was not significant in highly pathogenic avian influenza H7N9 and H5Nx datasets. This may indicate that these emerging viruses with avian-dominated host populations have a different immune selection pressure pattern compared to human influenza viruses.

**Figure 1.**
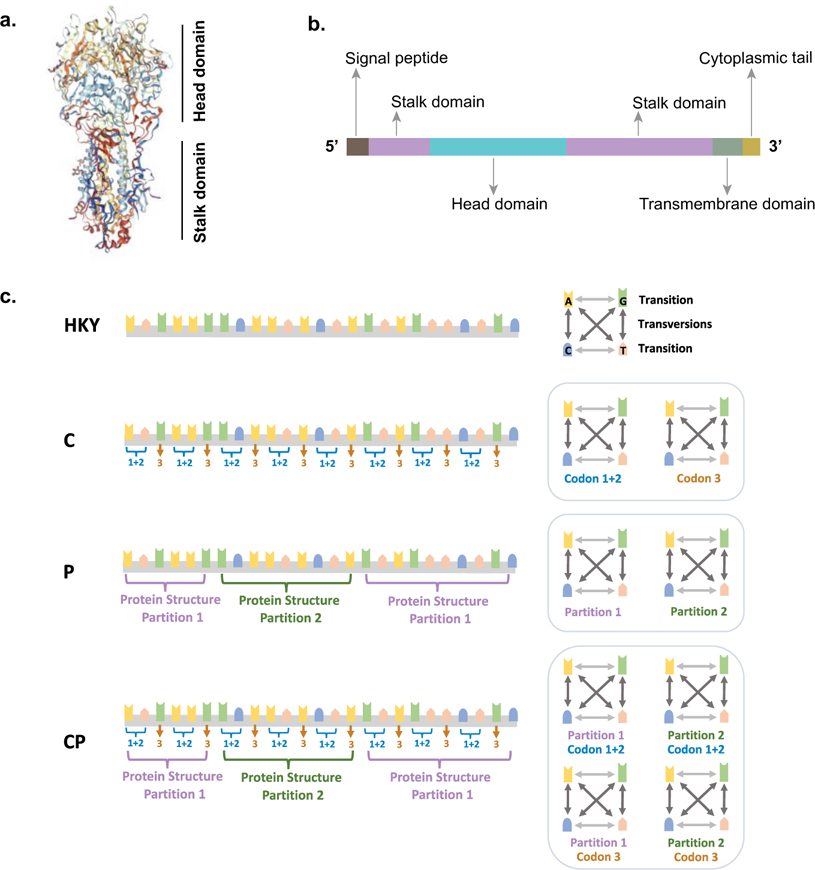
Conceptual diagram for influenza virus Hemagglutinins (HA) domains and four models tested ***1a. HA protein 3D structure: Stalk and Head domains.*** Head domain is the immunodominant with high plasticity, which contains the receptor binding sites. Stalk domain is immune-subdominant and more conserved. ***1b. Linear diagram for functional domains on HA nucleotide sequence***. The stalk domain has two parts separated by head domain on the linear sequence. To compensate for short length, signal peptide, transmembrane domain, and cytoplasmic tail are combined into one partition, referred to as “STC” in the analysis. ***1c. Model conceptual design.*** Four models are applied and compared for each dataset. **HKY model** is a substitution model that considers different base frequencies and assigns different rates for transitions v.s. transversions. **C model** represents SRD06 codon position model. The partitioning strategy is to analyze codon positions 1 + 2 and codon position 3 separately. **P model** takes the protein structure partitions into account based on the amino acid positions for each domain on the linear diagram. **CP model** combines both C and P models, which estimates codon positions in protein structural partitions. All models use HKY substitution model.

### Tree root height and substitution rates

To evaluate the accuracy and stability of model performance, tree parameters and estimated substitution rates were used to compare different models. Results from four models showed very similar tree root height and 95% Bayesian Credible Interval (BCI) for each influenza subtype (Table 1). The estimated mean substitution rates (Fig. 3) estimated from all models were similar for each dataset suggesting that the CP model generated reliable estimations for important evolutionary parameters that were comparable to other models. Furthermore, since both CP and P model incorporated partitioning scheme on protein structure, these two models provided domain-specific substitution rates for STC (including signal peptide, transmembrane domain and cytoplasmic tail), stalk and head domains, respectively. However, the domain-specific rates reported by P model were underestimated, compared to CP model (results shown in Supplemental Table 2). Such an observation was unsurprising since the HKY substitution model on each protein structure partition cannot account for more realistic variations among codon positions. One thing to note, though STC domain was not partitioned on codon positions, its small amount of information from short nucleotide length (about 150 nucleotides) resulted in high uncertainty of the STC substitution rates, showing long tails in the violin plots.

**Table 1.**
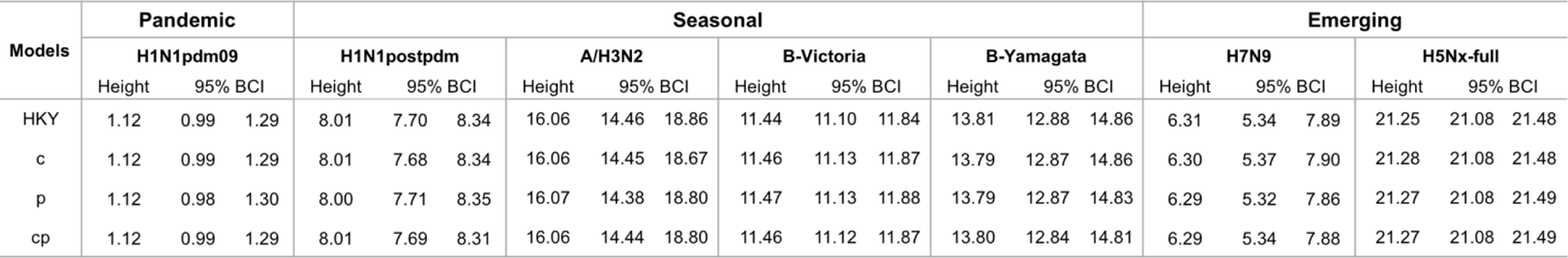
Tree root height estimation and its 95% Bayesian credible interval (BCI) from each model for each influenza subtype. Results from four models show very similar tree root height and 95% BCI for each influenza subtype. The unit of the root height is in years. **Models: HKY model** is a substitution model that considers different base frequencies and assigns different rates for transitions v.s. transversions. The **c model** represents SRD06 codon position model. The partitioning strategy is to analyze codon positions 1 + 2 and codon position 3 separately. The **p model** takes the protein structure partitions into account based on the amino acid positions for each domain on the linear diagram. The **cp model** combines both c and p models, which estimates codon positions in protein structural partitions. All models use HKY substitution model.

**Figure 2.**
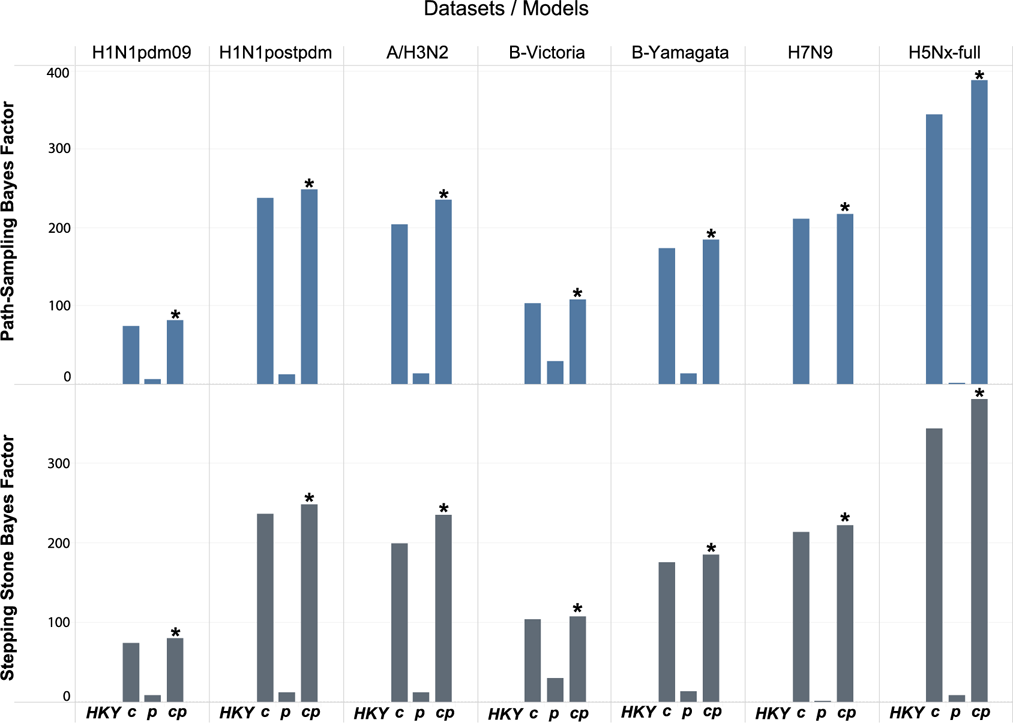
Log scale Bayes Factors for model selection procedure for each influenza subtype. All these Bayes Factors (BF) are calculated to compare with HKY via path-sampling (upper panel) and stepping-stone sampling (lower panel) approaches to compute the marginal likelihood estimates for each model. The asterisk * means a significant supportive BF for the structurally informed codon model (cp), compared to SRD06 codon only model (c model). Both cp and c models perform better than HKY model with decisive BF supports in all datasets. The HKY plus structurally informed model (p model) perform better in all datasets except for H7N9. Log scale BF criteria: BF< 5 represents no significance; 5 ≤ BF< 10 means substantial support; 10 ≤ BF< 15 means strong support; 15 ≤ BF < 20 means very strong support; BF ≥ 20 means decisive support. **Models: HKY model** is a substitution model that considers different base frequencies and assigns different rates for transitions v.s. transversions. The **c model** represents SRD06 codon position model. The partitioning strategy is to analyze codon positions 1 + 2 and codon position 3 separately. The **p model** takes the protein structure partitions into account based on the amino acid positions for each domain on the linear diagram. The **cp model** combines both c and p models, which estimates codon positions in protein structural partitions. All models use HKY substitution model.

**Figure 3.**
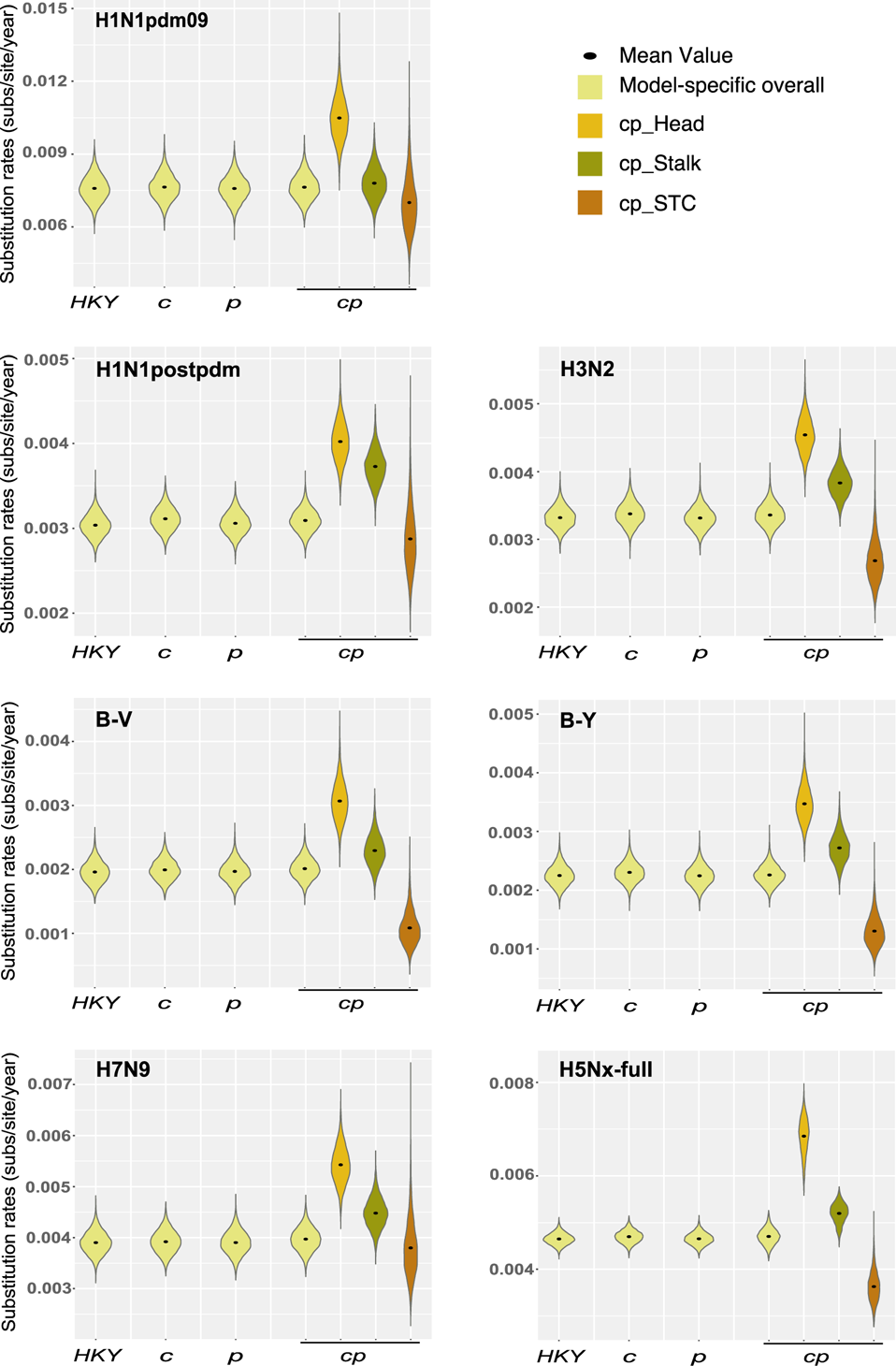
Overall mean substitution rates estimated from four models and domain-specific substitution rates from structurally informed partitioning model. Violin plots show the substitution rates from each step of Bayesian simulation. The results show overall mean substitution rates from four models for each dataset are similar, but the head domain has a significantly faster rate than that of stalk domain based on the new model (cp model). Generally, STC domain has more uncertain estimation with long tails of the violin plot, which is probably due to short length of nucleotides. Bayes Factors for the statistical tests on the differences of head and stalk domain substitution rates are 1,137 for H1N1pdm09, 6 for H1N1postpdm, 577 for A/H3N2, 197 for B-Victoria-like, 191 for B-Yamagata-like, 303 for H7N9 and +Infinity for H5Nx. **Models: HKY model** is a substitution model that considers different base frequencies and assigns different rates for transitions v.s. transversions. The **c model** represents SRD06 codon position model. The partitioning strategy is to analyze codon positions 1 + 2 and codon position 3 separately. The **p model** takes the protein structure partitions into account based on the amino acid positions for each domain on the linear diagram. The **cp model** combines both c and p models, which estimates codon positions in protein structural partitions. All models use HKY substitution model.

Comparing the overall substitution rates for each influenza subtype, we observed that the pandemic H1N1pdm09 had the highest substitution rate (∼7.5 × 10-3 substitution/site/year). HPAI H7N9 and H5Nx had the substitution rates around 4.0 – 4.8 × 10-3 substitution/site/year. Seasonal influenza viruses had the range as 2.0 – 3.5 × 10-3 substitution/site/year, where the two type B lineages had lower rates than the two A subtypes.

Furthermore, we compared the separate substitution rates for stalk and head domains from the CP model. The absolute rates calculated directly from each step of the Bayesian Markov Chain Monte Carlo (MCMC) simulation were plotted into violin plot (Fig. 3). Results showed that the head domain had a significantly higher substitution rate than stalk domain for all influenza subtypes, under the unit of substitutions per site per year. The Bayes Factors (BF) for the statistical tests between the substitution rates of head and stalk domains were 1137 (H1N1pdm09), 6 (H1Npostpdm), 577 (A/H3N2), 197 (B-Victoria), 191 (B-Yamagata), 303 (H7N9), +infinity (H5Nx), respectively, with the significant criteria as BF>3.

### Approximate selection pressure

With partitioning on codon positions, the CP model can estimate the ratio of substitution rate of codon positions 1 and 2 (majority of non-synonymous changes occur in these codons) over the rate of codon position 3 (majority changes are synonymous), that is, *d*C_1+2_/*d*C_3_, for stalk domain and head domain separately. The relative rate of C_3_ to C_1+2_ is a good predictor of the ω parameter of selection pressure measure (18) and thus could reflect the domain specific selection pressure. As shown in Table 2.2, the range of *d*C_1+2_/*d*C_3_ was 0.10 – 0.32 for stalk domain and 0.25 – 0.87 for head domain from all subtypes of influenza viruses included in the study. The overall *d*C_1+2_/*d*C_3_ for the whole HA protein calculated from the c model was lower than head domain but higher than stalk domain for each influenza subtype. We also observed that the stalk domain had a significantly lower *d*C_1+2_/*d*C_3_ than head domain for each influenza subtype (Bayes Factors shown in Table 2), which indicated that stalk domain experienced stronger purifying selection to maintain its conservation. Further to compare pandemic H1N1pdm09 and seasonal A/H1N1 post pandemic, we found that the seasonal H1N1 post pandemic had lower *d*C_1+2_/*d*C_3_ in both stalk and head domains, indicating pandemic H1N1pdm09 experienced less purifying selection.

**Table 2.** Substitution rate and dC1+2/dC3 of stalk and head domain for each influenza subtype. Compared to the head domain, the stalk domain with lower value of *d*C_1+2_/*d*C_3_ experiences stronger purifying selection to maintain its conserved functionality.

| Datasets | Substitution rates (subs/site/year) | | | | $dC_{1+2}/dC_3^*$ | | | |
| --- | --- | --- | --- | --- | --- | --- | --- | --- |
|  | Stalk 1+2* | Stalk 3** | Head 1+2 | Head 3 | Stalk | Head | BF<sub>++</sub> | Overall<sup>^</sup> |
| **H1N1pdm09** | 3.80E-03 | 1.18E-02 | 6.78E-03 | 1.42E-02 | 0.32 | 0.48 | 53 | 0.42 |
| **H1N1postpdm** | 1.26E-03 | 6.37E-03 | 2.16E-03 | 6.07E-03 | 0.20 | 0.36 | +Inf | 0.28 |
| **A/H3N2** | 1.39E-03 | 6.28E-03 | 3.66E-03 | 5.42E-03 | 0.22 | 0.68 | +Inf | 0.43 |
| **B-Victoria** | 5.44E-04 | 4.04E-03 | 1.51E-03 | 4.63E-03 | 0.13 | 0.33 | 8804 | 0.21 |
| **B-Yamagata** | 5.12E-04 | 4.93E-03 | 1.39E-03 | 5.56E-03 | 0.10 | 0.25 | 8875 | 0.18 |
| **H7N9** | 1.48E-03 | 7.48E-03 | 3.05E-03 | 7.80E-03 | 0.20 | 0.39 | +Inf | 0.30 |
| **H5Nx-full** | 1.74E-03 | 7.23E-03 | 6.11E-03 | 7.03E-03 | 0.24 | 0.87 | +Inf | 0.43 |

+: $dC_{1+2}/dC_3$ means the ratio of substitution rate of codon positions 1 and 2 over the rate of codon position 3.
++: BF is Bayes Factor. BF>3 represents statistical significance.
\*: 1+2 represent the mean substitution rates of the codon positions 1 and 2.
\*\* : 3 means the codon position 3.
<sup>^</sup>: overall $dC_{1+2}/dC_3$ is calculated from the SRD06 codon model for the whole HA gene.

To observe different patterns of selection pressure on the viral populations, phylogenetic tree branch-specific stalk *d*C_1+2_/*d*C_3_ and head *d*C_1+2_/*d*C_3_ were calculated and visualized via mapping the upper and lower quantiles of the ratio values onto the phylogenetic tree. We used seasonal H3N2 (Supplemental Fig. 1) and Egypt H5Nx (Supplemental Fig. 2) as examples. The *d*C_1+2_/*d*C_3_ had a range of 0.63 – 0.75 for H3N2 head domain, 0.21 – 0.23 for H3N2 stalk domain, 0.12 – 1.87 for Egypt H5Nx head domain and 0.09 – 0.32 for Egypt H5Nx stalk domain. Again, results from the branch-specific *d*C_1+2_/*d*C_3_ demonstrated that the head domain had significantly higher *d*C_1+2_/*d*C_3_ compared to stalk domain, which indicated that stalk domain was under stronger purifying selection pressure to maintain its conservation. To note, the majority of the *d*C_1+2_/*d*C_3_ values were less than 1, which indicated purifying selection, but occasionally some diversifying selection pressure (*d*C_1+2_/*d*C_3_ >1) was observed in Egypt H5Nx head domain in chicken-dominated population. This high *d*C_1+2_/*d*C_3_ in H5Nx chicken population may indicate that the viruses circulating in chicken may experience less purifying selection pressure to generate higher diversity with keeping more non-synonymous changes compared to that in human population. This suggests that avian population may generate more variants with pandemic potential.

### Sensitivity analysis and model validation

With the largest dataset HPAI H5Nx in this study, sensitivity analysis was performed to estimate the stability of model performance under different sample sizes. The same model fitting and selection procedure was tested on randomly selected 80%, 60%, 40% and 20% of the H5Nx-full dataset. The model selection procedure (Fig. 4) showed that all these datasets had decisive BFs to favor the structurally informed partitioning model with SRD06 (CP model). Tree root height estimation (Table 3) from these datasets had similar estimates and 95% BCI, where it was only slightly overestimated in the 20% of H5Nx dataset. The substitution rates of protein structure partitions (Fig. 5) also showed stable estimations across different sample sizes but only had slight overestimation with the 20% H5Nx dataset. Taken together, though the new model with partitioning on both codon positions and protein functional domains introduced more parameters, it performed stably with different amount of data and thus was not sensitive to sample size.

**Figure 4.**
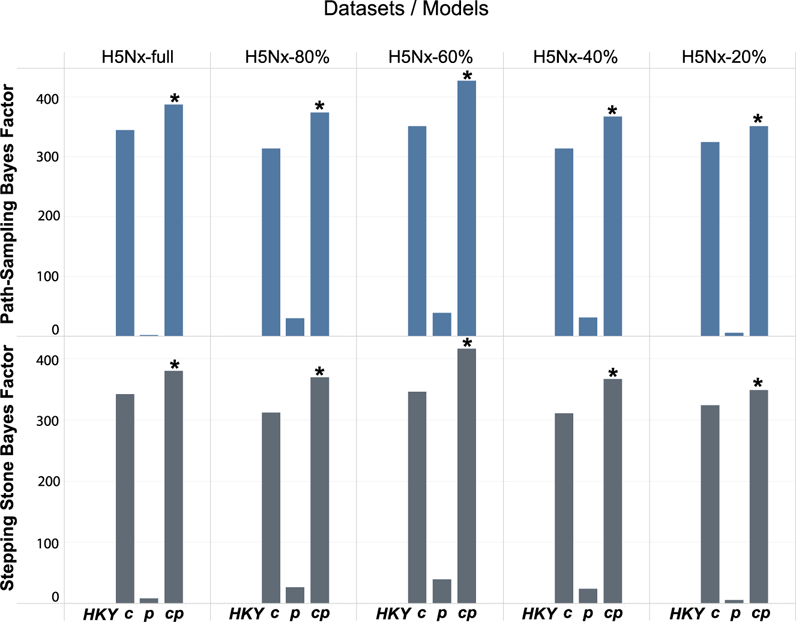
Log scale Bayes Factors for sensitivity analysis on full and subsampled HPAI H5Nx datasets. All these Bayes Factors (BF) are calculated to compare with HKY via path-sampling (upper panel) and stepping-stone sampling (lower panel) approaches to compute the marginal likelihood estimates for each model. The asterisk * represents a significant supportive BF for the structurally informed codon model (cp), compared to SRD06 codon only model (c model). Both cp and c models perform better than HKY model with decisive BF supports. The HKY plus structurally informed model (p model) performs better with supportive BFs in all datasets. Log scale BF criteria: BF< 5 represents no significance; 5 ≤ BF< 10 means substantial support; 10 ≤ BF< 15 means strong support; 15 ≤ BF < 20 means very strong support; BF ≥ 20 means decisive support. **Models: HKY model** is a substitution model that considers different base frequencies and assigns different rates for transitions v.s. transversions. The **c model** represents SRD06 codon position model. The partitioning strategy is to analyze codon positions 1 + 2 and codon position 3 separately. The **p model** takes the protein structure partitions into account based on the amino acid positions for each domain on the linear diagram. The **cp model** combines both c and p models, which estimates codon positions in protein structural partitions. All models use HKY substitution model.

**Figure 5.**
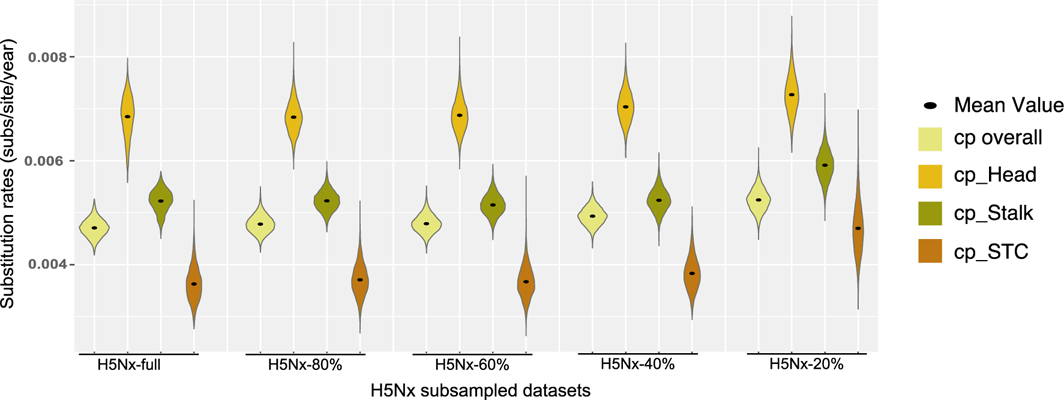
Sensitivity analysis: overall mean substitution rates and domain-specific substitution rates from structurally informed partitioning model for HPAI H5Nx subsampled datasets. Violin plots show the substitution rates from each step of the Bayesian simulations. Only 20% dataset has the trend to slightly exaggerate the rates and would not report the precise evolution history of H5Nx. The model superiority generally holds in the sensitivity analysis and the model is not sensitive to sample size. **Models:** The **cp model** combines both c and p models, which estimates codon positions in protein structural partitions. It uses HKY substitution model.

**Table 3.** Tree root height estimation and its 95% Bayesian credible interval (BCI) from each model for all H5Nx datasets. The statistical significance of the new model holds across different subsampling datasets. Only 20% dataset has the trend to slightly overestimate the root height and would not report the precise evolution history of H5Nx. In sum, the new model is not sensitive to sample size. The unit of root height is in years. **Models: HKY model** is a substitution model that considers different base frequencies and assigns different rates for transitions v.s. transversions. The **c model** represents SRD06 codon position model. The partitioning strategy is to analyze codon positions 1 + 2 and codon position 3 separately. The **p model** takes the protein structure partitions into account based on the amino acid positions for each domain on the linear diagram. The **cp model** combines both c and p models, which estimates codon positions in protein structural partitions. All models use HKY substitution model.

| Models | H5Nx-full |  |  | H5Nx-80% |  |  | H5Nx-60% |  |  | H5Nx-40% |  |  | H5Nx-20% |  |  |
| --- | --- | --- | --- | --- | --- | --- | --- | --- | --- | --- | --- | --- | --- | --- | --- |
|  | Height | 95% BCI |  | Height | 95% BCI |  | Height | 95% BCI |  | Height | 95% BCI |  | Height | 95% BCI |  |
| HKY | 21.27 | 21.08 | 21.48 | 21.31 | 21.08 | 21.57 | 21.34 | 21.07 | 21.62 | 21.42 | 21.07 | 21.79 | 21.64 | 21.07 | 22.19 |
| c | 21.29 | 21.08 | 21.48 | 21.30 | 21.08 | 21.57 | 21.32 | 21.07 | 21.61 | 21.43 | 21.07 | 21.81 | 21.65 | 21.08 | 22.20 |
| p | 21.28 | 21.08 | 21.49 | 21.32 | 21.08 | 21.57 | 21.32 | 21.07 | 21.59 | 21.41 | 21.07 | 21.79 | 21.65 | 21.08 | 22.20 |
| cp | 21.28 | 21.08 | 21.49 | 21.33 | 21.08 | 21.58 | 21.32 | 21.07 | 21.59 | 21.42 | 21.07 | 21.80 | 21.66 | 21.09 | 22.23 |

With two subsets of random selection from seasonal H3N2, model selection procedure and evolutionary parameter estimations generated very similar results. It indicated that the new model validated through different subsets and was not sensitive to the different distributions of epidemiological factors, including geographic location and isolation time (shown in Supplemental Table 3).

With dividing the H1N1pdm09 dataset into the Early stage and Later stage of an epidemic outbreak, model selection procedure reported that the CP model was significantly better than all other models in both datasets. The overall and domain-specific substitution rates in the Early stage was slightly higher compared to the Later stage of the outbreak (shown in Supplemental Table 4). And the Early stage had a higher *d*C_1+2_/*d*C_3_, indicating it experienced weaker purifying selection pressure to be relatively diverse. This may be related to host adaptation in the Early stage, where it generated more variants for the advantage of widely spreading in the host population.

## Discussion

In this study a novel phylogenetic model with integrating both codon positions and protein structure was developed and tested with HA protein of influenza viruses. Results showed that the new model was statistically supported to fit the data better and provided novel biological information that can reveal new insights into viral evolution. The substitution rates of stalk and head domains of HA protein were significantly different, with a higher rate in head domains for all the tested influenza types. Approximate selection pressure constructed from codon position partitions showed that stalk domain experienced higher purifying selection pressure to maintain its conservation. Model validation from subsets of multiple influenza subtype reported the model performed stably and was not sensitive to sample size, data distribution and epidemic stage. Taken together, this new model could provide potential biological explanations related to antigenic drift, vaccine escape and pandemic emergence to understand viral evolution and risk assessment.

Partitioning scheme on genetic data allows for capturing similar evolutionary features inside one partition of data and comparing different partitions to generate new insights on viral evolution. For example, SRD06 codon model categories the first and second codon positions together since they behave similarly to mainly generate non-synonymous changes, while the third codon position is treated as a separate partition due to mostly generating synonymous changes (3, 10). Some models conduct partitioning scheme on multiple proteins to reconstruct the co-evolution and estimate protein-specific rates under one model (19). These models function well for some purposes of observing the co-evolution of multiple proteins, however, it assumes evolutionary processes act uniformly across one protein, which may not be the biological reality due to distinctively functional domains on one protein.

When developing the new model in this study, the primary tasks are to efficiently capture the information regarding HA protein structural and functional domains and to properly incorporate this information into the evolutionary models. Therefore, how to partition on the protein structure with genetic data is critical. One solution is to compare all possible partitioning schemes for a given genetic sequence dataset to find the best partitioning strategy (3). However, this approach is computationally inflexible because the combination of possible partitioning schemes is tremendous even for a very small number of data blocks (20). Consequently, many previous studies ended up either arbitrarily choosing a single partitioning scheme or just selecting the best scheme based on statistical model selection (3,21,22). With these limitations on proper partitioning, the accuracy of the inferences from partitioned phylogenetic analysis remains uncertain, even though advanced developments in phylogenetic analysis have been achieved in recent decades (3). Due to advanced progress in viral protein structure (23), partitioning scheme in this study is based on biological functions of different protein domains, where HA protein has been defined to distinguish the immunogenic roles and other molecular-level functions during viral infection. Molecular mechanism of influenza viruses, for example, the initiation of viral fusion or antigenic-immune stimulation, is determined by the head domain of the protein. Nucleic acid changes observed in this domain usually result from a balance between immune selection and conserved functionality (24–26). The new partitioning model considers the distinctive functions of head domain to mainly bind to host-cell receptors and the stalk domain to perform membrane fusion.

Another advance in this study is that a formal model selection procedure with reliable approaches (27) to quantitatively evaluate the model performance and accuracy of inferences for pandemic, all seasonal and two emerging influenza viruses. Previously the harmonic mean estimate (HME) and Akaike’s information criterion through MCMC (AICM) (28) have been commonly used. In recent years, several new approaches to perform model selection in the field of phylogenetics, such as path-sampling (PS) (29) and stepping-stone sampling (SS) (30). Baele et al. (31) have tested HME, AICM, PS, and SS approaches for demographic and molecular clock model comparison, which “confirmed that HME systematically overestimates the marginal likelihood and fails to yield reliable model classification, and PS and SS substantially outperform HME estimators” (32). Therefore, in this study both PS and SS approaches have been used to perform the model selection procedure to generate reliable results of model comparison.

The new model with both partitioning on codon positions and protein structure incorporates biological functional domains inside one protein to allow for a quantitative exportation of the evolutionary process as a function of approximate host immune selection. The model did capture the different evolutionary process of these functional domains, which showed that the stalk domain had a significantly lower substitution rates and lower *d*C_1+2_/*d*C_3_, indicating that the stalk domain experienced stronger purifying selection and maintained its conserved functionality. The estimated domain-specific *d*C_1+2_/*d*C_3_ can reveal insights of selection pressure on different viral isolates in different host populations. For example, diversifying selection pressure on the head domain has been observed only in chicken population in Egypt. One possible explanation could be that the failed efficacy of poultry influenza H5 vaccines implemented in Egypt (33, 34) may drive the genetic diversity of the viruses in the chicken population.

Even though *d*C_1+2_/*d*C_3_ estimated by the model is not an exact measurement of the ω parameter, it is a good predictor of the ratio of non-synonymous changes over synonymous changes (10). The new model can provide an informative approximation and avoid heavy computational demands of a full codon or amino acid models (10). When assuming synonymous changes do not affect viral fitness, the *d*C_1+2_/*d*C_3_ value can provide corresponding estimation for selection pressure compared to mutation-selection models which assess the strength of natural selection on specific mutations (35). Furthermore, the partitioning scheme and additional parameter estimation did not significantly increase the analysis run time compared to other models in this study under the same computing environment.

The structurally informed evolutionary model may provide insight for universal vaccine design. The higher rates observed in the head domain are probably due to strong immune selection targeted on the globular head, where some sites are mostly targeted by the neutralizing immunity induced by seasonal influenza vaccine (13). Supported by many laboratory studies, stalk domain could be used to induce more broad-reactive vaccines (13,15,17). This model can more accurately reconstruct the ancestral sequence of the stalk domain with estimating domain-specific rates and substitution patterns under a shared viral phylogeny. Therefore, if this model was used to reconstruct common ancestral sequence for all viral types in influenza A virus Group 1 (H1, H2, H5, H6, H8, H9, H11, H12, H13, H16, and H17) and Group 2 (H3, H4, H7, H10, H14 and H15) (25), respectively, it is probably amongst the most promising approaches to design a universal influenza vaccine.

While this model has been developed with influenza in mind, there is potential to apply this approach to other viruses. Proteins, such as the respiratory syncytial virus F and G protein or the coronavirus spike protein that have multiple functional domains including immune stimulators, may be well suited to the novel approach that incorporates protein structural and functional domains into the nucleotide substitution models. Incorporating this type of information into comparative genomic analyses has the potential to provide important biological information and improve both vaccine design and evaluation.

## Methods

### Datasets

Multiple datasets representing pandemic, seasonal and emerging influenza viruses were used to develop and broadly test the proposed structurally informed partitioning models, including pandemic H1N1pdm09, seasonal influenza viruses A/H1N1postpdm, A/H3N2, B/Yamagata-like and B/Victoria-like, and two HPAI H5Nx and H7N9. Except for HPAI H7N9, H1N1pdm09 and seasonal A/H1N1postpdm that have been directly downloaded from the Global Initiative on Sharing All Influenza Data (GISAID, https://www.gisaid.org/) (36, 37), all other datasets were from published research, including seasonal A/H3N2 (38), HPAI H5Nx (39), B/Yamagata-like and B/Victoria-like (40). To avoid improper weights of duplicate isolates on the partitioning models, sequences with 100% similarity were removed for all datasets (final datasets can be found via the GitHub link: https://github.com/XuetingQiu/FluPartitioningModels).

Briefly, the final datasets contained 505 global isolates for H1N1pdm09 during 2009/04 – 2009/12, 635 global isolates for HPAI H7N9 during 2013-2018, and 554 global isolates for seasonal A/H1N1postpdm during 2012-2018. HPAI H5Nx contained 1,095 isolates during 1996-2016, after removing the vaccine candidates (39). The two influenza B virus lineages were full genome datasets sampled during 2002-2013 in eastern Australia and New Zealand (40). For B-Victoria, the dataset finally retained 214 isolates after randomly sampling down to 50% of the original dataset and removing the duplicates. For B-Yamagata, 241 isolates were included after removing the duplicates. The global dataset of seasonal A/H3N2 (38) contained 906 isolates during 2000-2012 after the duplicates were removed and further randomly subsampled.

### Definition of HA protein structural partitions

HA glycoprotein has been defined as different functional domains, including signal peptide, stalk domain, globular head domain, transmembrane domain and cytoplasmic domain (11, 12). These protein domains labeled with sites of amino acid were translated into the linear sites on the nucleotide sequences of HA gene (Fig. 2.1a and 2.1b). To use the full length of HA sequences and avoid some partitions being too short and lack of information, signal peptide, transmembrane domain, and cytoplasmic domain were combined as one partition named STC, given the similarly evolutionary characteristics of these functional domains. Though the stalk domain on the linear nucleotide sequence was separated by the head domain, the two gene regions of stalk domain were merged together for analysis (supplemental table I-1 and xml files in GitHub https://github.com/XuetingQiu/FluPartitioningModels).

### Structurally informed phylogenetic models

As shown in Fig. 2.1b, four different Bayesian phylogenetic models were conducted using the same molecular clock model and coalescent model for each dataset. Only the partitioning strategies on the nucleotide sequences were different. The base model applied HKY substitution model (4) with Gamma invariant distribution, where this model accounts for base frequencies and allows for rate variations between transitions and transversions. SRD06 codon model (C model in Fig. 2.1c) (10) partitioned all codons of the HA gene reading frame into codon positions 1+2 and codon position 3, where these two partitions also used HKY substitution model. Protein structure partitioning model (P model in Fig. 2.1c) contained STC, stalk and head functional domains with HKY substitution model for each domain. The last model (CP model in Fig. 2.1c) combined the partitioning strategy in both C and P models, that is, codon positions in each protein functional domain were further partitioned into codon positions 1 + 2 and codon position 3.

### Tree likelihood construction in the partitioning models

After partitioning with codon positions or/and protein structural domains, the overall tree likelihood calculations need to be explained. The general Bayesian inferences are based on the posterior probability of a hypothesis, for example, a given tree 𝜏. Then the posterior probability of the tree based on the data can be obtained using Bayes theorem,

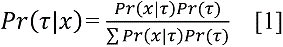

In which, *Pr(*𝜏) is the prior probability of the tree hypothesis 𝜏 based on data x; *Pr(x|*𝜏) is the probability of observing the data at a specific site.

Then assuming independence of substitution across sites, the general likelihood based on a specific phylogenetic model, i.e. the probability of observing the aligned matrix X of all j sequences is,

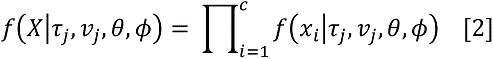

In which, c is the number of total sites along the sequences; 𝜏 represents the tree hypothesis, 𝑣 represents the branch lengths, 𝜃 is the substitution model parameters, and 𝜙 represents all other model parameters.

The proposed structurally informed partitioning model based on HA protein functional domains extends the formula [2] of the tree likelihood estimation. Each of the defined partition has its tree likelihood estimated independently, therefore, the overall tree likelihood is the products of each partition-specific likelihood, that is,

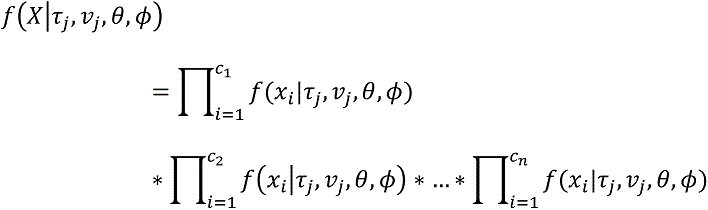

In which, the sum of c_1_, c_2_, …, c_n_ is equal to c, the total number of nucleotide sites along the sequences; and n is the number of partitions in the model.

### Phylogenetic model simulations

All models were simulated in the package of Bayesian Evolutionary Analysis Sampling Trees (BEAST) v1.8.4 (41). To allow for rate variations across lineages, the uncorrelated lognormal relaxed molecular clock was used with an initial mean of 0.0033 with a uniform prior ranging from 0.0 to 1.0. Based on prior knowledge, a smooth and time-aware Gaussian Markov random field (GMRF) process prior on the population sizes was applied in the Skyride calescent model (42). Each model was performed at least four independent runs of 100 million Markov Chain Monte Carlo (MCMC) generations. To report the substitution rates and phylogenetic trees, four runs for each model were combined after removal of burn-in to achieve an Effective Sample Size (ESS) of >200 as diagnosed in Tracer v 1.5. Visualized violin plots for substitution rates were generated via personal R scripts (GitHub https://github.com/XuetingQiu/FluPartitioningModels).

### Model selection procedure and criteria

The goal of model selection procedure is to compare model superiority and identify a model that is sufficiently complex to capture the biological realism and evolutionary processes that have occurred but to avoid overparameterized models with more parameters than these can be reliably estimated from the available data (3, 31). In this study, both path-sampling (PS) (29) and stepping-stone sampling (SS) (30) were used to compute the marginal likelihood estimation to perform the model selection procedure. PS and SS approaches account for both the number of parameters and the appropriateness of prior distributions for these parameters.

The marginal likelihood is the probability of the data (that is, likelihood) given the model type, not assuming any particular model parameters (31). The marginal likelihood estimation for both PS and SS with Beta path step distributions is set to run 100 path steps and length of chains as 1 million. Bayes Factors (BF), the ratio of marginal likelihood estimations from two models, are used to evaluate the significance of the model comparison. When converted to a log scale, BF becomes the difference between two log marginal likelihood estimations. The mathematical conversion is below (43):

For a model with parameters Θ, the marginal likelihood for the model M with data X is,

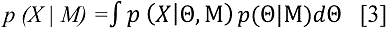

Then the Bayes factor can be calculated for model M1 against model M2,

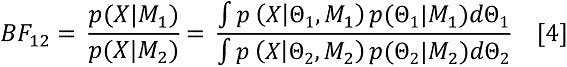

The log scale of BF is derived as below,

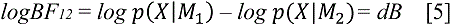

With log scale BF as dB, the criteria for model selection are: dB <0 means the selection supports M2; 0 ≤ dB <5 means no evidence for supporting M1; 5 ≤ dB < 10 means substantial strength of evidence for supporting M1; 10 ≤ dB <15 means strong evidence of supporting M1; 15 ≤ dB < 20 means very strong evidence of supporting M1; dB ≥ 20 means decisive evidence of supporting M1.

### Model performance, validation and sensitivity analysis

To compare model performance among four models for each dataset, two basic evolutionary parameters were estimated and reported: the root heights and overall substitution rates. If the parameters for each dataset were similar across these four models, then it indicated that the model can provide accurate and stable estimation no matter how many partitions it has included.

To evaluate whether the structurally informed partitioning model was sensitive to sample size, sensitivity analysis was performed on HPAI H5Nx from 1996-2017, the largest final dataset with 1,095 isolates in this study. With retaining the oldest isolate of H5Nx, the rest was randomly subsampled to keep 80%, 60%, 40% and 20% of the dataset, respectively. Four runs of 100 million MCMC chain length of the four models for the full dataset and these subsampled datasets were conducted with marginal likelihood estimations for model selection. Tree height parameters, domain-specific substitution rates, and model selection BFs were reported to evaluate the data sufficiency for model stability.

To further conduct model validation, two more approaches were applied. Firstly, to evaluate whether the structurally informed partitioning model was sensitive to data distribution, the full seasonal A/H3N2 dataset was randomly and repeated sampled 40% of the full dataset into two subsets. These two subsets had different geographical and temporal distributions of A/H3N2 isolates. Secondly, to assess whether the new model was sensitive to the epidemic stage. With the H1N1pdm09 dataset, the Early stage of the pandemic was defined as April to middle of July, 2009, and the Later stage of the pandemic was July-December, 2009. The same model variations and model selection procedure were conducted to all these subsets.

### Domain-specific d*C_1+2_/*d*C_3_* recorded to tree branches

The new model can report domain-specific rates and *d*C_1+2_/*d*C_3_ values. To visualize these specific rates or ratios on the tree branches, the BEAST xml files (GitHub link: https://github.com/XuetingQiu/FluPartitioningModels) were coded to compute and load the specific branch rates for stalk domain codon positions 1+2, stalk domain codon position 3, head domain codon positions 1+2, and head domain codon position 3. Python script (GitHub link: https://github.com/XuetingQiu/FluPartitioningModels) was constructed to compute and write the *d*C_1+2_/*d*C_3_ for stalk and head domain from the median rate of each partition to the final summarized tree branches. Seasonal A/H3N2 and all Egypt H5Nx data were used as examples to show the domain-specific *d*C_1+2_/*d*C_3_ changes in A/H3N2 with different vaccine isolates introduced and the changes in different hosts (human v.s. avian) in Egypt H5Nx.

## Acknowledgements

This study was partially funded by NIAID Centers of Excellence for Influenza Virus Research and Surveillance (CEIRS, HHSN272201400006C). These funders had no role in study design, data collection and analysis, decision to publish, or preparation of the manuscript. We gratefully acknowledge the authors, originating and submitting laboratories of the sequences from GISAID’s EpiFluTM Database, for making tremendous data publicly available and largely benefiting the science community.

