## SupplementalTablesFigures for "Structurally informed evolutionary models improve phylogenetic reconstruction for emerging, seasonal, and pandemic influenza viruses"

**Supplemental Table 1. The nucleotide sites of functional partitions for each influenza subtype.** The nucleotide sites are corresponding to the aligned HA open reading frame of nucleotide sequences for each influenza subtype. STC represents the combination of signal peptide, transmembrane domain and cytoplasmic tail. HPAI means highly pathogenic avian influenza.

| Influenza subtypes | STC domain | Stalk domain | Head domain |
|---|---|---|---|
| A/H1N1pdm09 | 1-51, 1591-1701 | 52-216, 874-1590 | 217-873 |
| A/H1N1postpdm | 1-51, 1591-1701 | 52-216, 874-1590 | 217-873 |
| A/H3N2 | 1-48, 1588-1701 | 49-204, 877-1587 | 205-876 |
| B-Victoria | 1-45, 1645-1758 | 46-171, 922-1644 | 172-921 |
| B-Yamagata | 1-45, 1642-1755 | 46-171, 919-1641 | 172-918 |
| HPAI H7N9 | 1-54, 1573-1683 | 55-216, 856-1572 | 217-855 |
| HPAI H5Nx | 1-48, 1597-1713 | 49-174, 868-1596 | 175-867 |

**Supplemental Table 2. The domain-specific rates from p model compared to cp model for each influenza subtype.** The results showed that p-model slightly underestimates the domain-specific rate compared to cp model. HPAI means highly pathogenic avian influenza.

**Models:** The **P model** takes the protein structure partitions into account based on the amino acid positions for each domain on the linear diagram. The **CP model** combines both c and p models, which estimates codon positions in protein structural partitions. Both models use HKY substitution model.

| Datasets | Substitution rates (subs/site/year) | | | |
|---|---|---|---|---|
| | P model | | CP model | |
| | Head | Stalk | Head | Stalk |
| **A/H1N1pdm09** | **9.18E-03** | **6.46E-03** | **1.05E-02** | **7.81E-03** |
| **A/H1N1postpdm** | **3.37E-03** | **2.98E-03** | **4.12E-03** | **3.81E-03** |
| **A/H3N2** | **3.87E-03** | **3.02E-03** | **4.54E-03** | **3.83E-03** |
| **B-Victoria** | **2.50E-03** | **1.68E-03** | **3.07E-03** | **2.29E-03** |
| **B-Yamagata** | **1.80E-03** | **7.73E-04** | **3.47E-03** | **2.72E-03** |
| **HPAI H7N9** | **4.45E-03** | **3.52E-03** | **5.43E-03** | **4.48E-03** |
| **HPAI H5Nx-full** | **5.26E-03** | **3.80E-03** | **6.85E-03** | **5.26E-03** |

**Supplemental Table 3. Model selection and parameter estimations from two subsets of H3N2.** The full H3N2 dataset is randomly sampled down to 40% to generate two subsets, which results in each containing 365 isolates from different geographical and temporal distributions. The results show that the new model performs stably for different subsets and are not sensitive the different distributions of the dataset by year and by country (Distributions of the two subsets are shown in Table I-3 in appendix figures a and b following the table).

**Models: HKY model** is a substitution model that considers different base frequencies and assigns different rates for transitions v.s. transversions. The **c model** represents SRD06 codon position model. The partitioning strategy is to analyze codon positions 1 + 2 and codon position 3 separately. The **p model** takes the protein structure partitions into account based on the amino acid positions for each domain on the linear diagram. The **cp model** combines both c and p models, which estimates codon positions in protein structural partitions. Both models use HKY substitution model.

| Parameters | H3N2 subset 1 | H3N2 subset 2 |
|---|---|---|
| **Bayes Factors by path-sampling** for model selection | | |
| HKY model | - | - |
| c model | 146.3 | 130.9 |
| p model | 5.2 | 6.0 |
| cp model | 167.3 | 144.5 |
| **Bayes Factors by stepping-stone** for model selection | | |
| HKY model | - | - |
| c model | 146.2 | 130.8 |
| p model | 5.2 | 5.7 |
| cp model | 167.8 | 144.0 |
| **Substitution rates** for each model | | |
| HKY model | 4.39E-03 [3.9596E-3,4.8401E-3] | 4.33E-03 [3.9024E-3,4.7817E-3] |
| c model | 4.51E-03 [4.0855E-3,4.9475E-3] | 4.43E-03 [3.9946E-3,4.8712E-3] |
| p model | 4.37E-03 [3.9456E-3,4.8208E-3] | 4.31E-03 [3.8896E-3,4.7522E-3] |
| cp model | 4.50E-03 [4.0775E-3,4.9391E-3] | 4.36E-03 [3.9300E-3,4.8035E-3] |
| **Root Height** for each model | | |
| HKY model | 13.851 [13.2406,14.7147] | 13.866 [13.2242,14.7571] |
| c model | 13.855 [13.2584,14.7190] | 13.881 [13.1729,14.7161] |
| p model | 13.858 [13.2862,14.7688] | 13.878 [13.2330,14.7818] |
| cp model | 13.859 [13.2574,14.7566] | 13.874 [13.1963,14.7553] |
| **Partition-Specific substitution rates** | | |
| Stalk domain | 5.08E-03 | 4.93E-03 |
| Head domain | 6.01E-03 | 5.71E-03 |

**Table 3 appendix figure a. Distribution of H3N2 subset 1 by year and by country.**

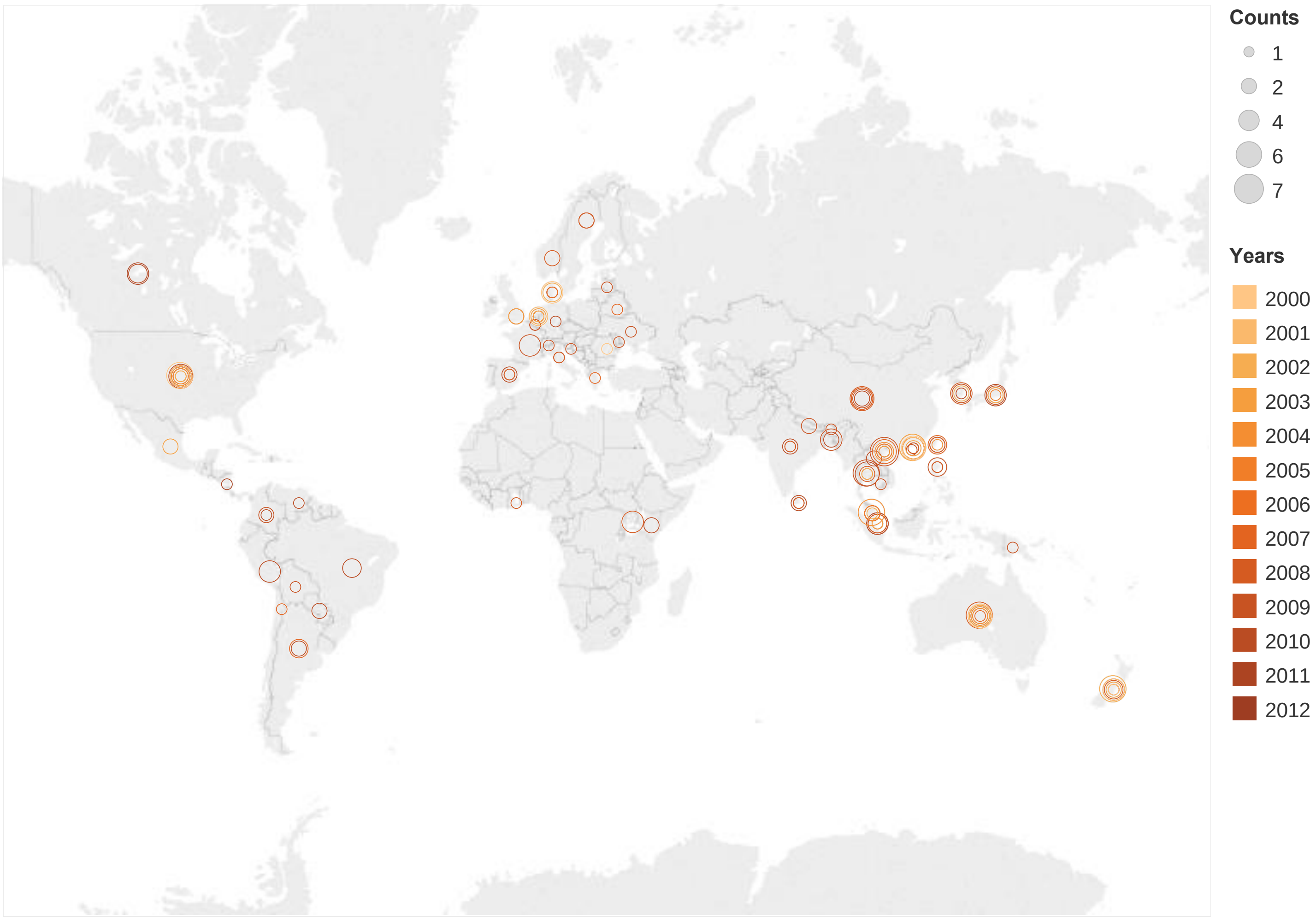

**Table 3 appendix figure b. Distribution of H3N2 subset 2 by year and by country.**

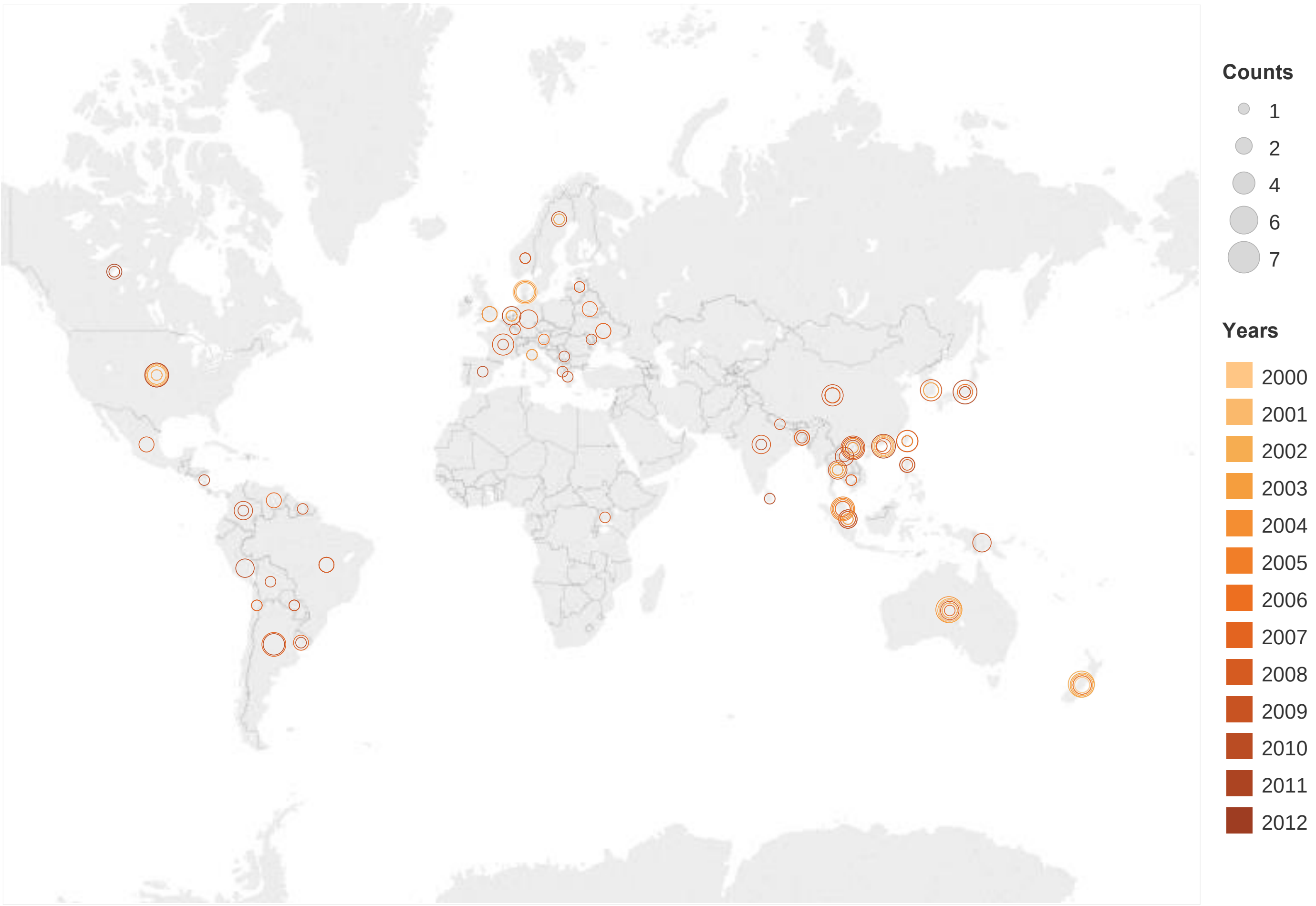

**Supplemental Table 4. Model selection and parameter estimation from the Early and Later epidemic stage of H1N1pdm09**. The early epidemic stage of H1N1pdm09 is defined as 04/01/2019-07/15/2019 with 274 isolates, and the Later epidemic stage of H1N1pdm09 is defined as 07/16/2019-12/31/2019 with 231 isolates. The results show that the superiority of the new model holds (BF>5 compared to HKY model) and performs stably (overall substitution rates, root height and domain-specific substitution rates are similar from each model) for dataset from different stage of epidemics. The substitution rates and overall $dC_{1+2}/dC_3$ are lower at the later stage of the pandemic than the early stage of the pandemic.

| Parameters | H1N1pdm09-Early | H1N1pdm09-Later |
|---|---|---|
| **Bayes Factors by path-sampling** for model selection | | |
| HKY model | - | - |
| c model | 34.0 | 28.2 |
| p model | 5.3 | 7.5 |
| cp model | 40.9 | 35.7 |
| **Bayes Factors by stepping-stone** for model selection | | |
| HKY model | - | - |
| c model | 33.6 | 27.5 |
| p model | 5.7 | 7.0 |
| cp model | 42.5 | 34.9 |
| **Substitution rates** for each model | | |
| HKY model | 9.66E-03 [7.54E-03, 1.20E-02] | 8.03E-03 [6.57E-03, 9.54E-03] |
| c model | 9.84E-03 [7.71E-03, 1.21E-02] | 8.06E-03 [6.65E-03, 9.54E-03] |
| p model | 9.66E-03 [7.49E-03, 1.19E-02] | 8.02E-03 [6.50E-03, 9.52E-03] |
| cp model | 9.75E-03 [7.51E-03, 1.20E-02] | 8.09E-03 [6.66E-03, 9.59E-03] |
| **Root Height** for each model | | |
| HKY model | 0.551 [0.4311, 0.7001] | 0.8801 [0.7322, 1.0806] |
| c model | 0.546 [0.4313, 0.7009] | 0.8821 [0.7353, 1.0773] |
| p model | 0.547 [0.4305, 0.6950] | 0.8819 [0.7379, 1.0863] |
| cp model | 0.551 [0.4334, 0.7066] | 0.8808 [0.7313, 1.0704] |
| **Partition-Specific substitution rates** | | |

| | | |
|---|---|---|
| Stalk domain | 9.36E-03 | 8.59E-03 |
| Head domain | 1.32E-02 | 1.10E-02 |
| **Overall $dC_{1+2}/dC_3$** | 0.45 | 0.39 |

**Supplemental Figure 1. H3N2 Branch Specific $dC_{1+2}/dC_3$ for Head and Stalk domain separately.** $dC_{1+2}/dC_3$ means the ratio of substitution rate of codon positions 1 and 2 over the rate of codon position 3. Figure a is for Head domain and b is for Stalk domain. These branch specific $dC_{1+2}/dC_3$ is the median value for each branch from the MCMC steps. The branch stroke weight is proportional to the value of $dC_{1+2}/dC_3$ for each tree. The blue branch color represents the branch specific $dC_{1+2}/dC_3$ is lower than the overall median of $dC_{1+2}/dC_3$ for all isolates, while the orange branch color means the branch specific value is higher than the overall median. Vertical lines indicate the introduction time of vaccine strains selected by WHO. Solid vertical line represents North Hemisphere vaccine strain and dashed line represents South Hemisphere vaccine strain. There were 13 vaccine strains introduced during a 10-year period.
Head domain has much higher $dC_{1+2}/dC_3$ (0.63-0.75) compared to stalk domain (0.21-0.23) but all $dC_{1+2}/dC_3$ are less than 1, which means the stalk domain is under higher purifying selection to maintain its conserved functionality. Some branches have higher $dC_{1+2}/dC_3$, where the color is orange and branch weight is thicker. Taken together, this model could provide potential biological and quantitative information to understand viral evolution.

a. Head domain

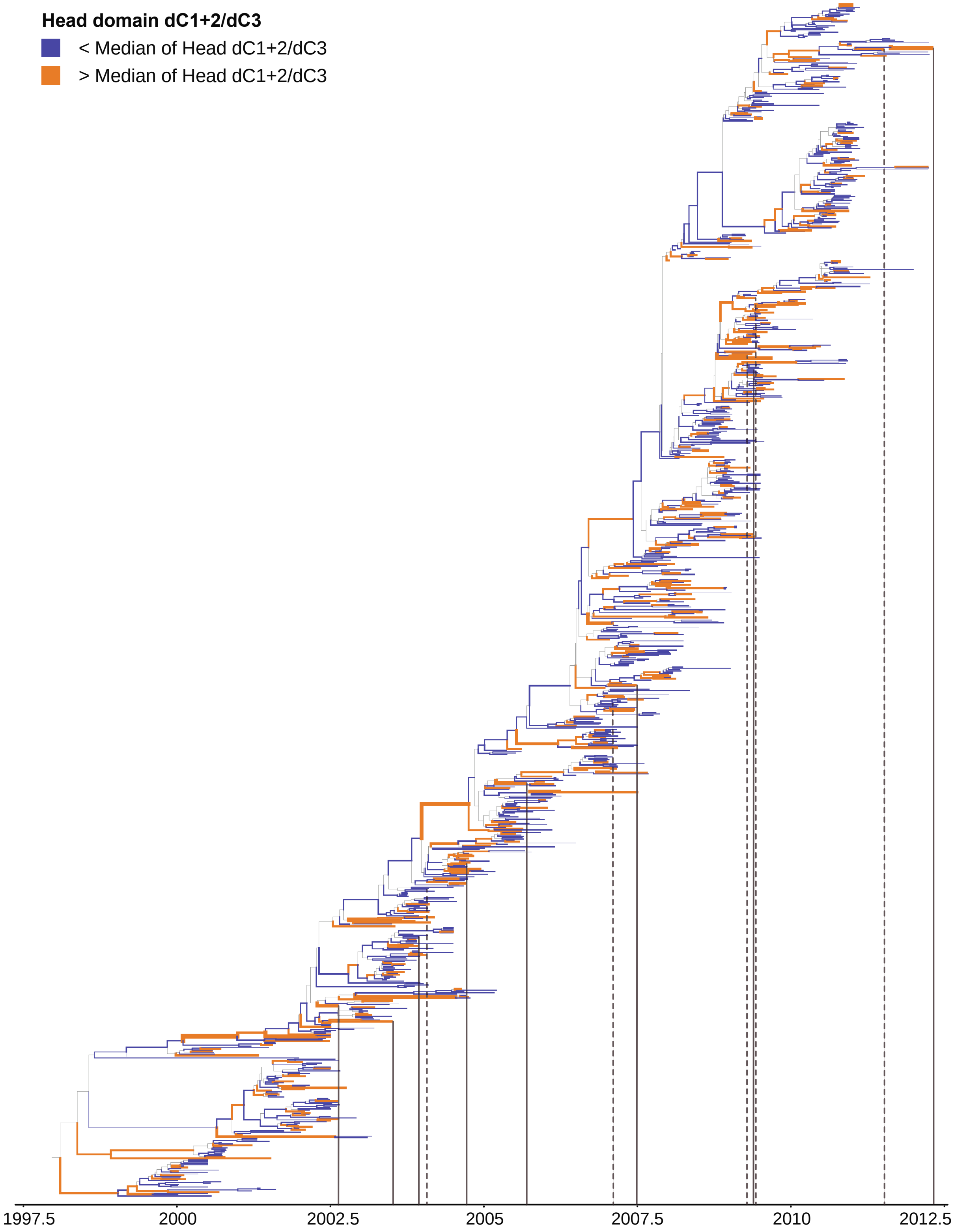

b. Stalk domain

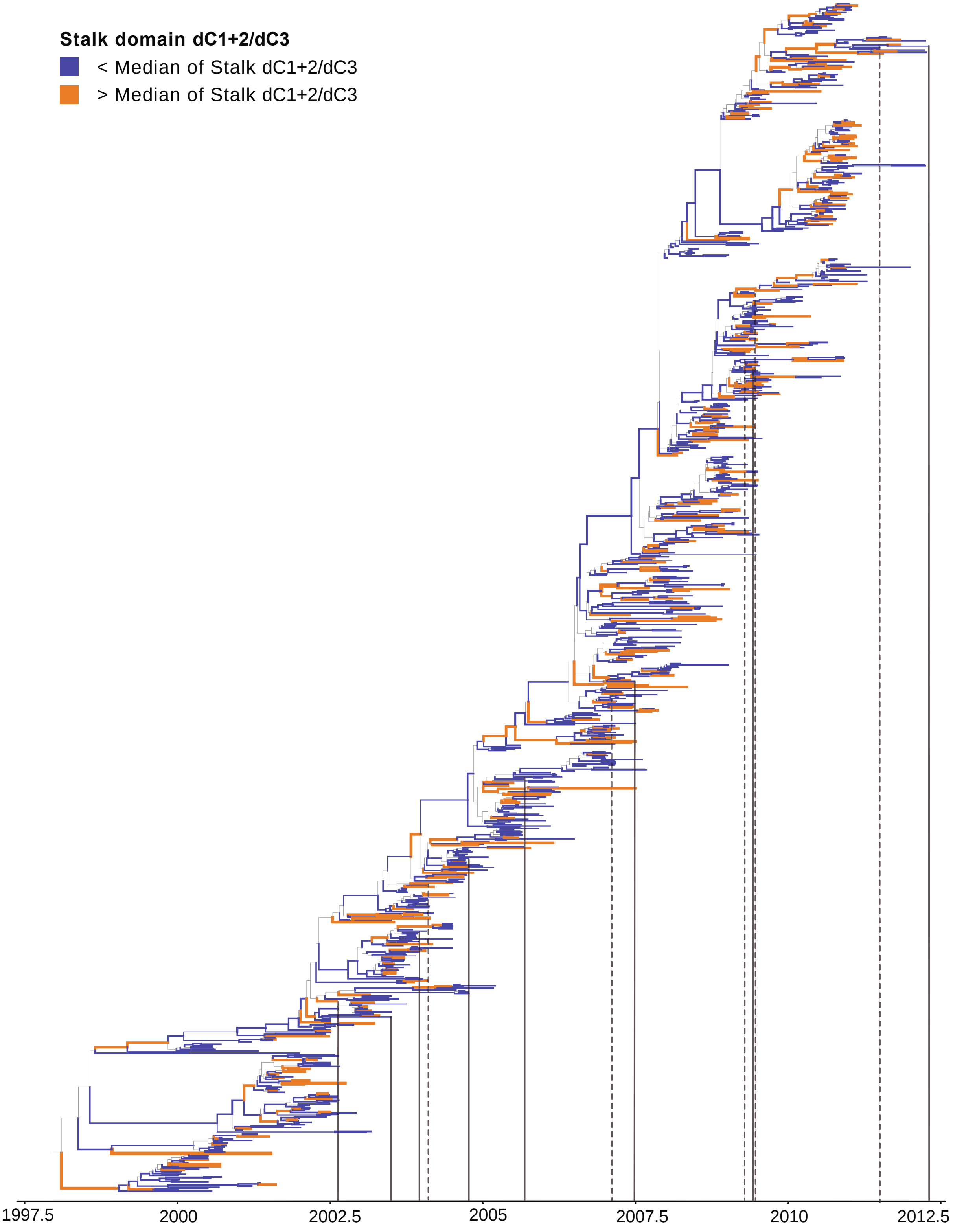

**Supplemental Figure 2. Egypt H5Nx Branch Specific $dC_{1+2}/dC_3$ for Head and Stalk domain separately.** $dC_{1+2}/dC_3$ means the ratio of substitution rate of codon positions 1 and 2 over the rate of codon position 3. Figure a is for Head domain and b is for Stalk domain. These branch specific $dC_{1+2}/dC_3$ is the median value for each branch from the MCMC steps. The branch stroke weight is proportional to the value of $dC_{1+2}/dC_3$ for each tree. The blue branch color represents the branch specific $dC_{1+2}/dC_3$ is lower than the overall median of $dC_{1+2}/dC_3$ for all isolates, while the orange branch color means the branch specific value is higher than the overall median. Head domain has much higher $dC_{1+2}/dC_3$ (0.12 – 1.87) compared to stalk domain (0.09 – 0.32), which means the stalk domain is under higher purifying selection to maintain its conservation and head domain occasionally experiences diversifying selection with $dC_{1+2}/dC_3$ >1 in avian population. Furthermore, higher $dC_{1+2}/dC_3$ are observed in avian viral branches, which means that viruses experience lesser purifying selection pressure to generate more diverse on non-synonymous changes in avian populations. Taken together, this model could provide potential biological explanations of host factors on viral evolution.

a. Head domain

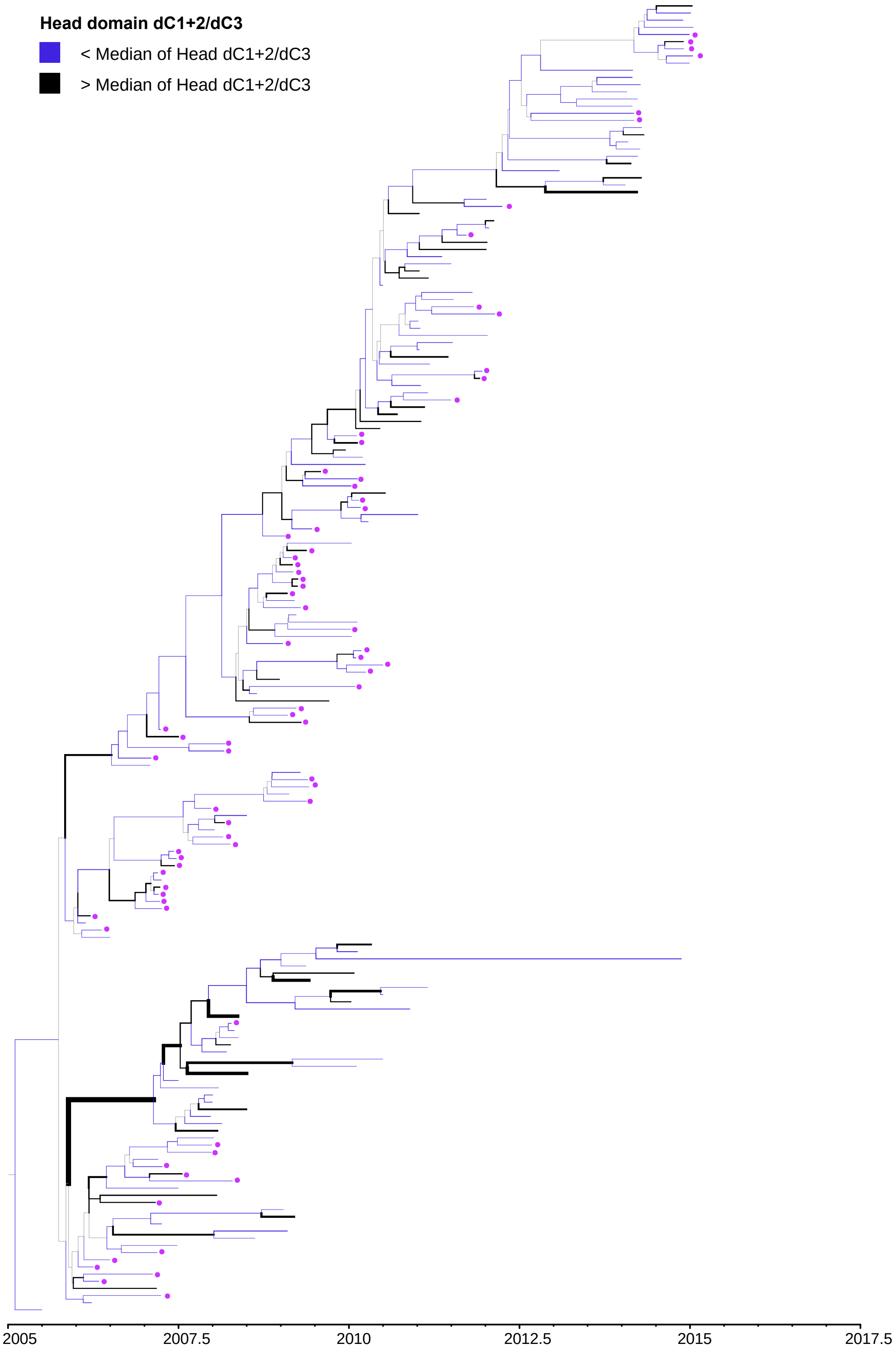

b. Stalk domain

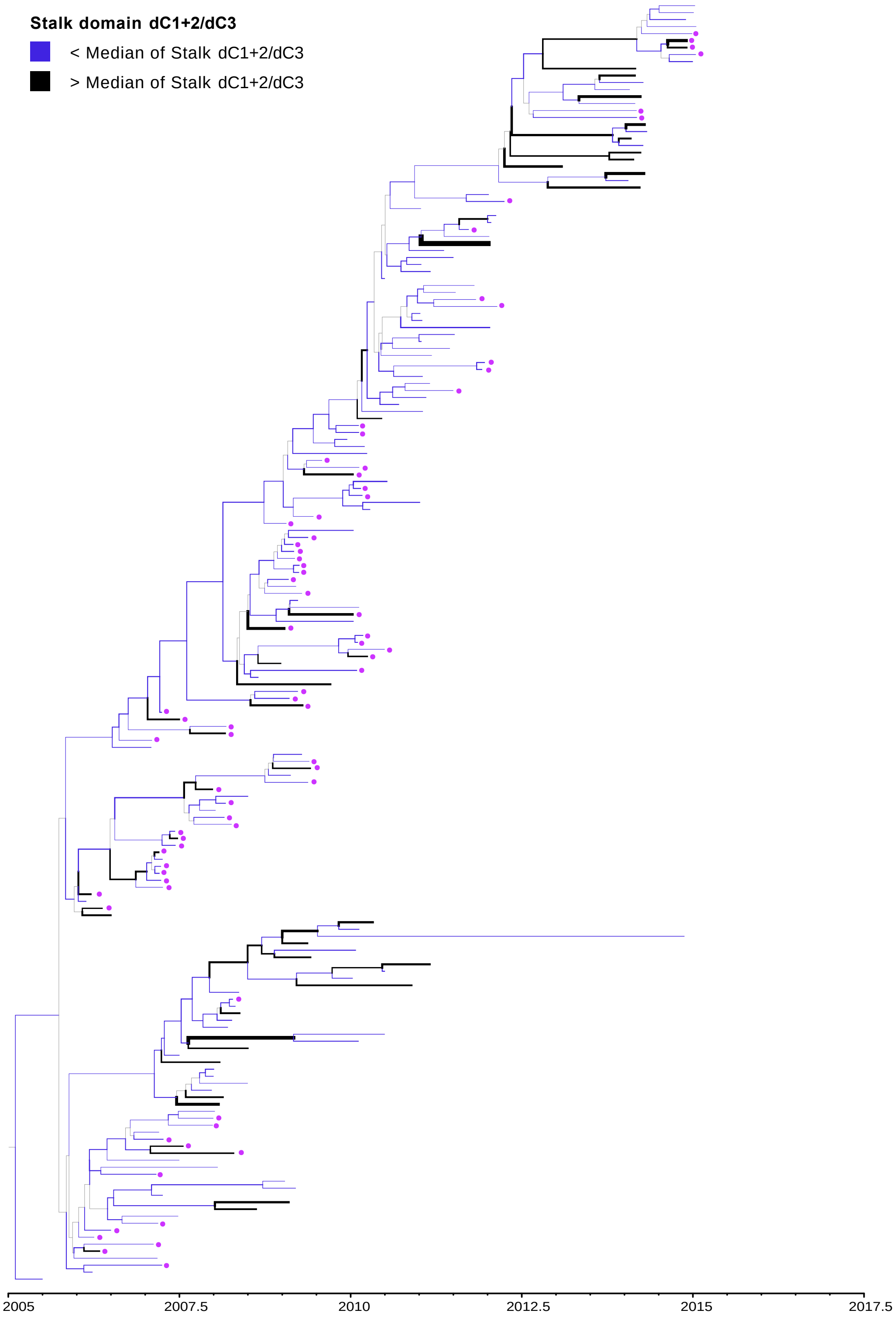
